# Identification of small molecules with virus growth enhancement properties

**DOI:** 10.1101/2022.11.08.515589

**Authors:** Ma Jesús García-Murria, Laura Gadea-Salom, Sandra Moreno, Oscar Zaragoza, Alejandro Brun, Ismael Mingarro, Luis Martínez-Gil

## Abstract

The novel severe acute respiratory syndrome coronavirus-2 (SARS-CoV-2) has caused the pandemic disease known as coronavirus disease 2019 (COVID-19). COVID-19 vaccines were developed at record speed and were authorized approximately a year after the original outbreak. This fast response saved the lives of countless individuals and reduced the disease burden of many more. The experience has served as a reminder of the necessity to implement solid vaccine development platforms and fast production pipelines. Manufacturing vaccines for enveloped viruses, including some SARS-CoV-2 vaccines, often relies on the production of large quantities of viruses in vitro. Thus, speeding up or increasing virus production would expedite vaccine development. With this objective in mind, we established a high throughput screening (HTS) to identify small molecules that enhance or speed up host-virus membrane fusion. Among the HTS hits, we identified that ethynylestradiol augments SARS-CoV-2 fusion activity in both the absence and presence of TMPRSS2. Furthermore, we confirmed that ethynylestradiol can boost the growth of not only SARS-CoV-2 but also Influenza A virus in vitro. A small molecule with these characteristics could be implemented to improve vaccines production.

**Importance:** The (COVID-19) pandemic had a tremendous impact on our healthcare systems and the global economy. The rapid development of effective vaccines saved the lives of countless individuals and reduced the disease burden of many more. Intending to increase vaccine production, we developed and performed a high-throughput screening (HTS) to identify small molecules that enhance viral and cellular membrane fusion. Among the HTS hits, we confirmed that Ethynylestradiol can boost the growth of SARS-CoV-2 and Influenza A virus *in vitro*.

## Introduction

The novel severe acute respiratory syndrome coronavirus-2 (SARS-CoV-2) (1,2) has caused the pandemic disease known as coronavirus disease 2019 (COVID-19). This disease has had and continues to have, a tremendous impact on our healthcare systems and the global economy. Accordingly, extraordinary efforts have been taken to develop prophylactic and therapeutic measures that can reduce the morbidity and mortality associated with COVID-19 disease. COVID-19 vaccines were developed at record speed and were authorized for use approximately a year after the original outbreak (3,4). This fast response saved the lives of countless individuals and reduced the disease burden of many more. The experience has served as a reminder of the necessity to implement solid vaccine development platforms and fast production pipelines.

The cellular membrane is the first barrier that the virus encounters when infecting a new cell. At some point during entry enveloped viruses must fuse the cellular and viral membranes. In the case of SARS-CoV-2, the highly glycosylated spike (S) protein is the one responsible for gaining entry to the host cells (5). The S protein, a class I trimeric fusion protein (6), is translated in a non-active form (S_0_). Proteolytic activation of S_0_ by host proteases produces the mature pre-fusion S protein that is incorporated in the virions (7). The infectious process starts with the binding of mature pre-fusion S to the cellular receptor Angiotensin-converting enzyme 2 (ACE2) at the cell surface (2,8). Binding to the receptor triggers a conformational change, resulting in the exposure of the fusion peptide which will interact with the host membrane and initiate the fusion of the viral and cellular membranes. Facilitating any of the multiple steps in this complex process would speed up and/or increase viral production.

The production vaccines for enveloped viruses, including some for SARS-CoV-2, often relies on the generation of large quantities of viruses *in vitro (9)*. Speeding up or increasing virus production would expedite vaccine development, reduce production costs and increase manufacture yield. With this objective in mind, we established a high throughput screening (HTS) to identify small molecules that enhance or speed up the growth of enveloped viruses *in vitro*. Among the HTS hits, we confirmed that ethynylestradiol can boost the growth of SARS-CoV-2 and Influenza A virus *in vitro*.

## Results and Discussion

### Implementation of the BiMuC approach

The novel SARS-CoV-2 is a highly contagious agent. Accordingly, its handling requires substantial safety measures which may complicate research protocols. To facilitate the study of the SARS-CoV-2 S-mediated membrane fusion process, we decided to implement a fluorescence-based complementation assay (BiMuC) (10). The BiMuC approach is a safe, virus-free approach based on the structural properties of some fluorescent proteins such as the Venus Fluorescence Protein (VFP) (11–13). Briefly, the VFP can be split into two fragments (VN and VC respectively), none of which is fluorescent by itself. However, if these two fragments are fused to a pair of interacting proteins which bring them together the native structure of the VFP is reconstituted and its fluorescence properties are recovered (Figure 1A). The aforementioned chimeras are expressed in two separate cell pools together with the viral machinery required for membrane fusion. If membrane fusion brings the fluorescent chimeras nearby the fluorescence would be reconstituted. On the contrary, no signal would be retrieved if the conditions for the formation of a syncytium are not met (Figure 1B).

**Figure 1.**
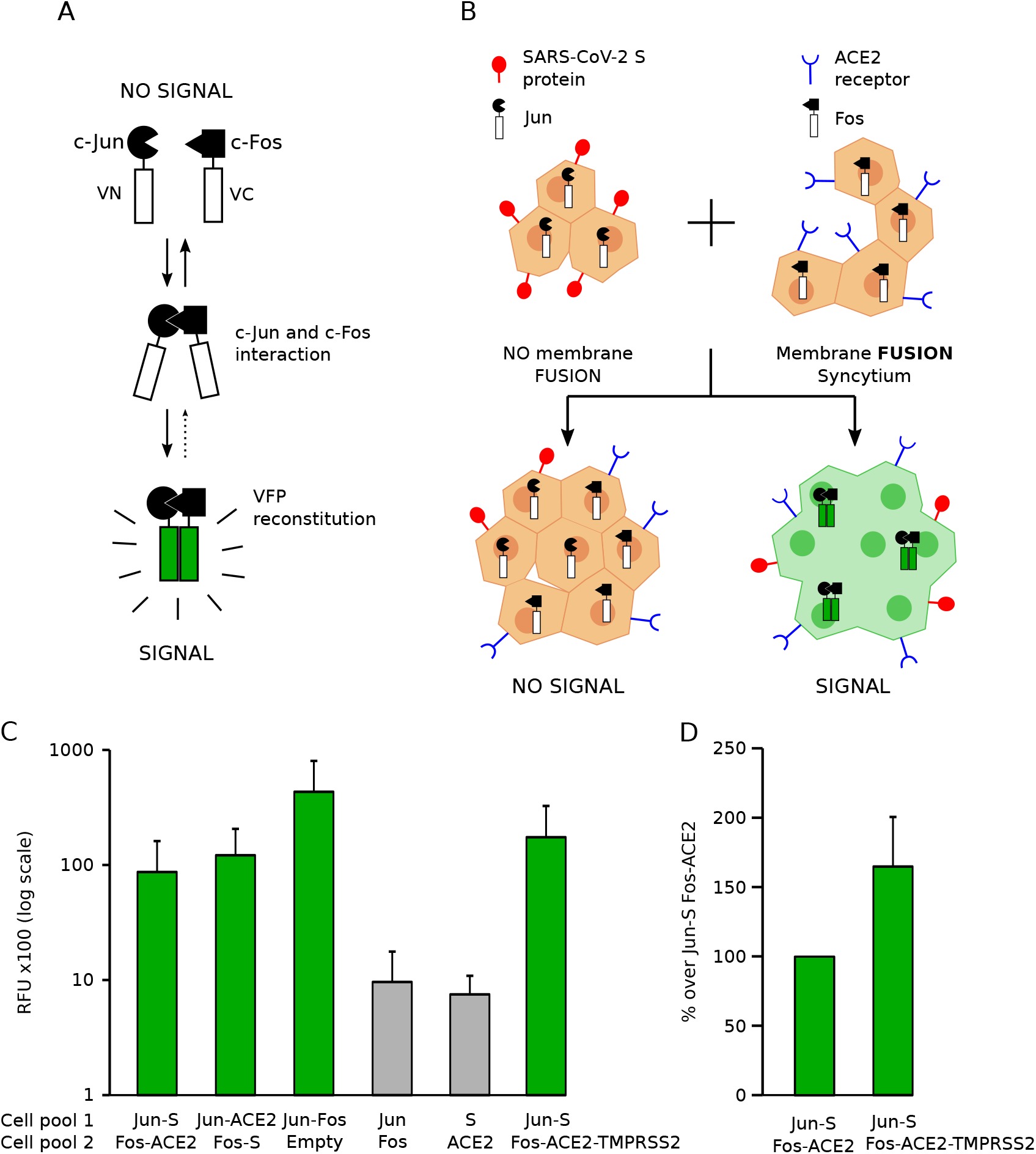
Analysis of SARS-CoV-2 membrane fusion by BiMuC. **A**. Schematic representation of the BiFC assay. The c-Jun and c-Fos proteins were fused to the Nt and Ct ends of the VFP, respectively, to create the c-Jun/VN and c-Fos/VC chimeras (Jun and Fos). Reconstitution of the structure of the VFP (represented in green), and thus its fluorescence properties, occurs exclusively after c-Jun and c-Fos interaction. **B**. Schematic representation of the BiMuC assay. To study SARS-CoV-2 membrane fusion properties. Hek-293T cells were transfected either with SARS-CoV-2 S and Jun or with ACE2 and Fos. Next, these two cell populations were combined. Successful interaction between SARS-CoV-2 S and ACE2 leads to the fusion of the cellular membranes, the creation of a syncytium, and the reconstitution of the VFP native structure (depicted in green). On the contrary, inhibition of any of the cellular or viral processes that lead to membrane fusion (from synthesis and maturation of the S protein to its interaction with ACE2) will result in the absence of a fluorescent signal. **C**. For the cell-cell fusion assay, Hek-293T cells with the indicated plasmid combinations. The next day cells were counted, mixed, and seeded into 96 well plates. After 48 hrs, fluorescence was measured. Bars show the average and standard deviation of at least three independent experiments. Green bars denote those samples with fluorescence levels comparable to the positive control (Jun-Fos). **D**. To facilitate visualization of the TMPRSS2 effect on viral fusion RFUs were plotted as the percentage over the Jun-S Fos-ACE2 sample.

To test whether we could apply this methodology to the study of SARS-CoV-2-induced membrane fusion the interacting partners c-Jun and c-Fos (14) were linked to the VN and VC fragments of the VFP, generating the c-Jun/VN and c-Fos/VC chimeras, from now on presented simply as Jun and Fos. These chimeras were expressed in two independent cell pools together with SARS-CoV-2 S (Jun-S) and ACE2 (Fos-ACE2) respectively. Additionally, we included cells transfected with Jun and Fos together (Jun-Fos), Jun, Fos, S, ACE2, or an empty plasmid (Empty) in the assay. After transfection, cell pools were combined as indicated (Figure 1C) and incubated for additional 48 hrs. The samples containing cells transfected with Jun-S and Fos-ACE2 showed a fluorescence signal significantly stronger than those that combined pools of cells expressing Jun and Fos or S and ACE2 (negative controls) suggesting that a successful fusion of the cell membranes triggered by the viral machinery brought the two chimeras together and facilitated reconstitution of the fluorescent signal. Note that we also tested the Jun-ACE2 and Fos-S combinations. This data indicates that the BiMuC methodology could be implemented to safely study SARS-CoV-2 mediated membrane fusion.

Our assay is based on Hek-293T cells, which express low levels of ACE2 and TMPRSS2 (15,16). We did not include TMPRSS2 in our assay to set up a sub-optimal fusion state that facilitates the identification of small molecules that enhance syncytia formation. To confirm that our assay can identify an increase in viral fusion we included TMPRSS2 in the transfection mix together with ACE2 (Fos-ACE2-TMPRSS2). The addition of the serine protease increased the observed fluorescent signal (Figure 1C and D).

### High throughput small molecule identification

In light of these results, we decided to perform a small molecule screening to identify compounds that could enhance membrane fusion. We adapted our methodology for the HTS and verified its suitability by the Z′ factor (0.64 or 0.62 for the Jun-S + Fos-ACE2 or Jun-ACE2 + Fos-S combinations respectively) (17). For the HTS, Hek-293T cells were transfected with Jun-S or Fos-ACE2. After transfection cell pools were mixed as described previously, seeded at 96 well plates (10^5^ cells/well), and treated with the appropriate small molecule. Each plate included a column of wells treated with DMSO. Additionally, plates included 8 wells as a negative control containing cells that had been transfected with Jun or Fos (both cell pools were combined as previously) and treated with DMSO. Once treated, cells were incubated for 48 hours followed by fluorescence measurement and hit selection.

A library consisting of 1280 chemically diverse small molecules (Prestwick Chemical, GreenPharma) was screened and the results were standardized by the *Z-Score;* primary hits were selected based on a *Z-Score* > 1.5. Next, hit compounds were repurchased from a different vendor and re-tested at multiple concentrations to confirm their ability to enhance viral membrane fusion (Figure 2A). Alongside, we assayed the toxicity of the compounds. Next, to identify whether the selected compounds could induce an increase in viral fusion under optimal growing conditions, we then tested them in the presence of TMPRSS2. In this case, the enhancing effect of the compounds was even stronger than without the protease (Figure 2B). In all assays [DMSO] was kept at ≤ 1 %.

**Figure 2.**
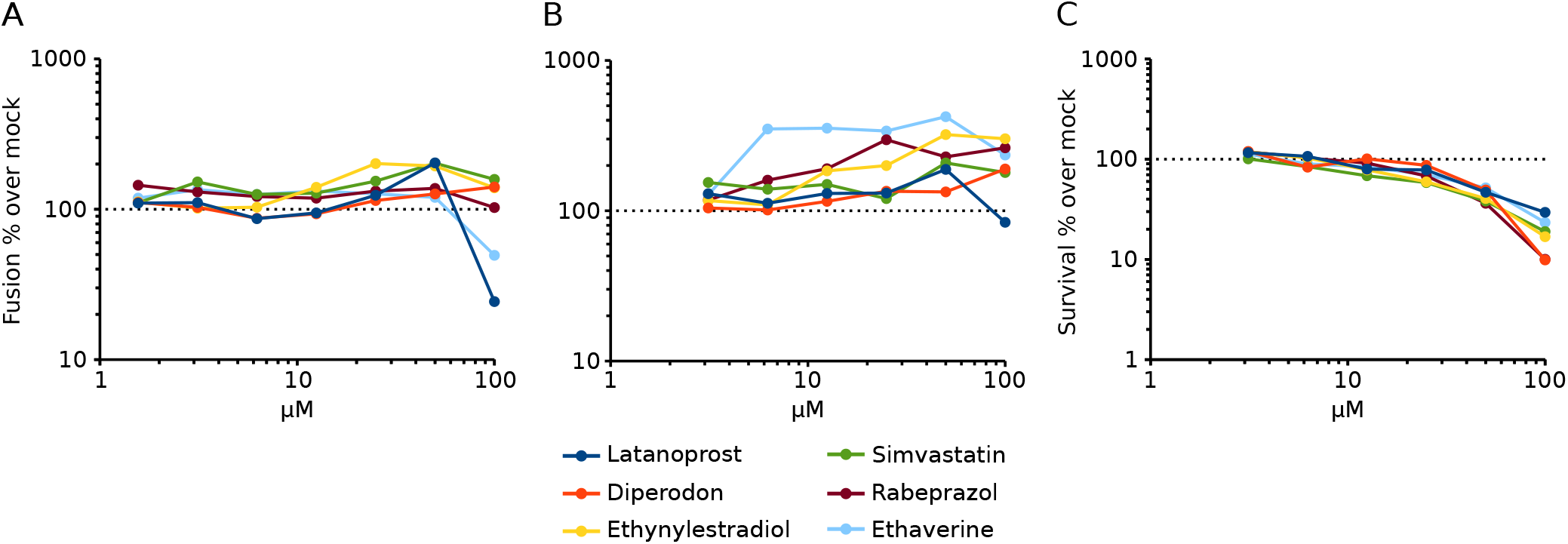
Analysis of hit compounds. **A**. and **B**. Selected compounds were repurchased and their ability to increase syncytia formation was tested with the BiMuC assay at the indicated concentration in the absence (A) or presence (B) of TMPRSS2. The fluorescence percentages over the mock (DMSO) treated samples are represented. Dots represent the average and standard deviation of at least three independent experiments. **C**. Toxicity was analyzed at the indicated concentration using the Cell-Titer Glo assay (Promega) following the manufacturer’s indications.

### Effect of Ethaverine, Rabeprazole, and Ethynylestradiol on the growth of SARS-CoV-2 and Influenza A virus

Our assay is based on syncytia formation, which might not recapitulate accurately the fusion of the viral and cellular membranes occurring during an infection. Consequently, at this point, we decide to analyze the effect of the hit compounds on the viral growth of SARS-CoV-2 (15,18,19). Based on our previous results we decided to focus on Ethaverine, Rabeprazole, and Ethynylestradiol. Our results indicated that Rabeprazole and Ethynylestradiol increased viral titers at 48 hrs post-infection (Figure 3A and B). However, Ethaverine did not increase or accelerate the growth of SARS-CoV-2 (Figure 3C).

**Figure 3.**
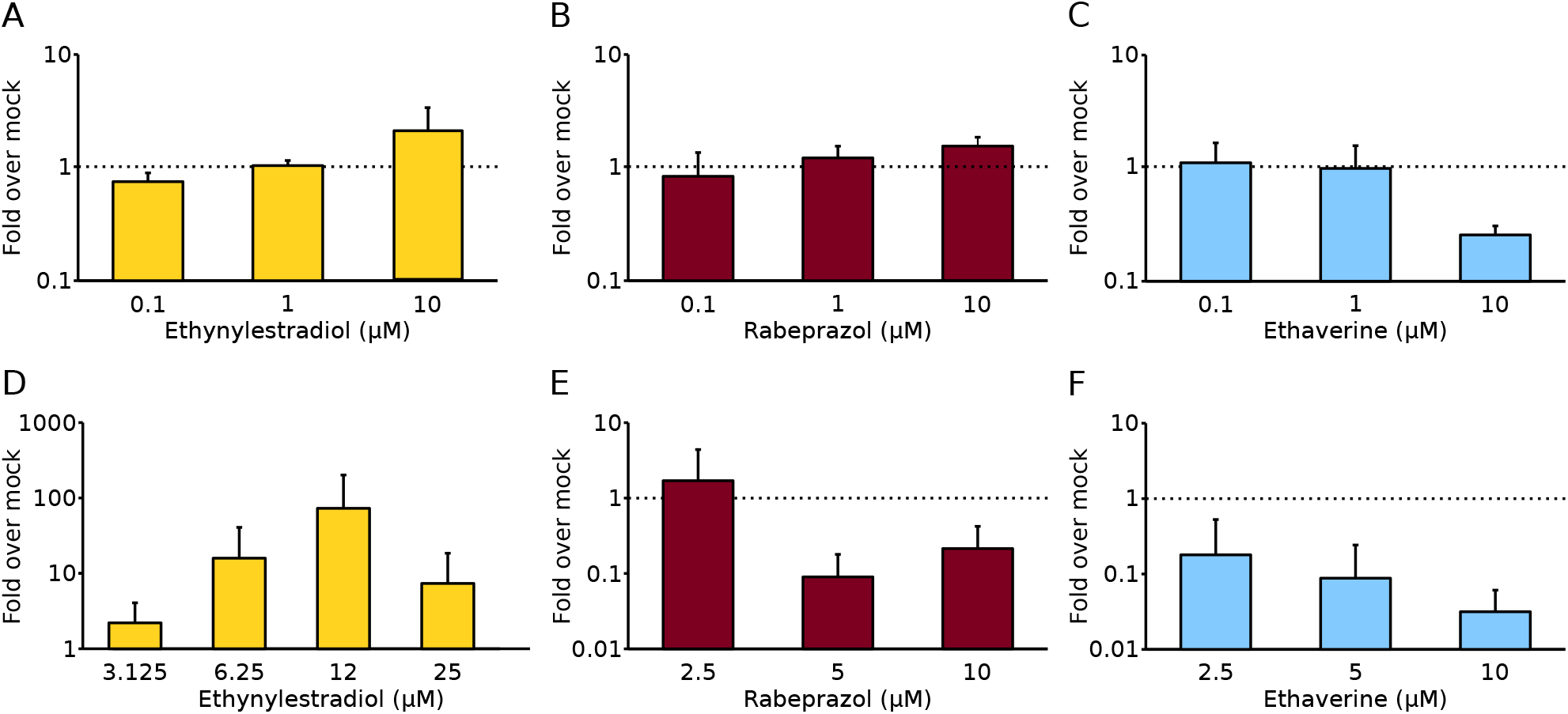
Effect of selected compounds on SARS-CoV-2 and IAV. **A to C**. SARS-CoV-2 was grown at an MOI of 0.01 in VeroE6 cells for 48h in the absence or presence of Ethaverine, Rabeprazole, or Ethynylestradiol at the indicated concentrations. Next, supernatants were collected and titered by standard plaque assay in VeroE6 cells. Bars show the average and standard deviation of at least three independent experiments. **E to C**. IAV/WSN/33, at an M.O.I of 0.001, was used to infect Madin-Darby canine kidney (MDCK) cells. IAV was incubated in the absence or presence of Ethaverine, Rabeprazole, or Ethynylestradiol at the indicated concentrations for 48h. Next, supernatants were collected and titered by the 50 % Tissue Culture Infective Dose (TCID50) method in MDCK cells. Bars show the average and standard deviation of at least three independent experiments.

Next, we analyzed if the ability of these compounds to enhance viral growth was restricted to SARS-CoV-2. To do so, we tested the effect of Ethaverine, Rabeprazole, and Ethynylestradiol on the growth of the Influenza A virus (IAV). IAV infections were carried out in the presence and absence of the selected compounds as previously described (20). Ethynylestradiol increased IAV titers by approximately 1 log (Figure 3D). On the other hand, Ethaverine and Rabeprazole did not increase viral titers (Figure 3E and F).

Most of the compounds included in the chemical library used in our HTS are approved drugs. However, it is important to note that our experiments are not sufficient to claim that the hit compounds, including Ethynylestradiol (Figure 4), might worsen the outcome of a patient during infection with an enveloped virus.

**Figure 4.**
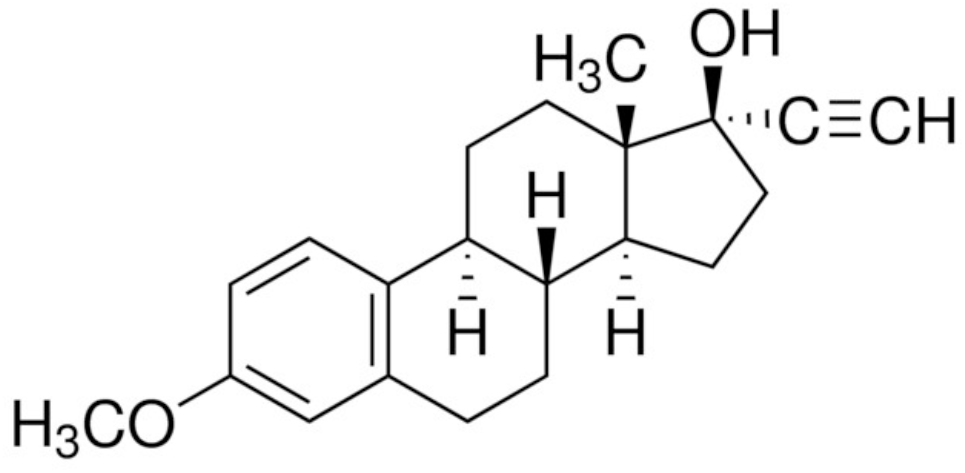
Structure of Ethynylestradiol. Structure of Ethynylestradiol 3-methyl ether. CAS#72-33-3. Smiles: [C@]12([C@]([C@]3([C@@](c4c(CC3)cc(cc4)OC)(CC1)[H])[H])(CC[C@]2(C#C)O)[H])C.

In summary, our results indicate that Ethynylestradiol could be used *in vitro* to enhance the growth of enveloped viruses. Based on previous works (21,22,22–25) we believe Ethynylestradiol might be altering the properties of the viral and/or cellular membranes to facilitate membrane fusion and or viral endocytosis. Compounds such as polybrene are used to improve retrovirus infections by facilitating the adhesion of virions to the cell surface, since the addition of positively charged polycations reduces the repulsive forces between the cell and the virus, improving transduction efficiency (26). However, as far as we are aware no compounds that improve membrane fusion and subsequently increase and accelerate viral growth had been identified previously. A small molecule with this characteristic could be easily implemented to increase production and/or reduce production times and costs.

## Materials and Methods

### Cell lines and plasmids

Hek-293T, VeroE6, and Madin-Darby canine kidney (MDCK) cells were obtained from ATCC (http://www.atcc.org) and were maintained in Dulbecco’s Modified Eagle Medium (DMEM) (Gibco, http://www.lifetechnologies.com) supplemented with 10 % fetal bovine serum (FBS, Gibco). The SARS-CoV-2 S plasmid was a gift from Dr. F. Krammer (Ichan School of Medicine at Mount Sinai). The Jun-Nt VFP (Jun) and Fos-Ct VFP (Fos) and the human ACE2 and TMPRSS2 expressing plasmids were obtained from Addgene (#22012, #22013, 1786, and #53887 respectively).

### BiMuc Assay

To study SARS-CoV-2 membrane fusion properties Hek-293T cells (DMEM supplemented with 10 % FBS) were transfected, using polyethyleneimine (PEI) (27), either with SARS-CoV-2 S and Jun or with ACE2 and Fos plus the pRL Renilla Luciferase (Promega) to normalize the signal. Additionally, we tested the SARS-CoV-2 S - Fos and the ACE2 – Jun combinations, once again with the pRL Renilla Luciferase. The next day cells were counted, mixed, and seeded into 96 well plates (3 × 10^4^ cells/well in 100 μL of media). After 48 hours, fluorescence was measured on 96 well plates as previously described (11).

For the high throughput identification of small molecules with the ability to modulate membrane fusion, the aforementioned protocol was adjusted to meet the necessary HTS standards. To evaluate the robustness of our assay we calculated the Z′-factor and the Signal-to-Noise (S/N) ratio. Z′-factor = 1 − ((3δpos + 3δneg)/(μpos − μneg)), where μpos is the mean signal for the positive control, μneg is the mean signal for the negative control, δpos is the standard deviation of the positive control, and δneg is the standard deviation for the negative control. S/N = (μpos − μneg)/((δpos) × 2 + (δneg) × 2) × 0.5.

Briefly, Kek-293T cells were seeded in 15 cm dishes (5 × 10^6^ cells/dish) using DMEM supplemented with 10% FBS. After 24 h (day 2) of incubation (37 °C, 5% CO _2_), cells were transfected using PEI with the aforementioned plasmid combinations or mock transfected. On the next day, cells were counted, mixed, and seeded into 96 well plates (3 × 10^4^ cells/well in 100 μL of media). Cells were mixed into 2 pools: Pool A containing cells expressing Jun-Nt and SARS-CoV-2 S or Fos-Ct and ACE2 and pool B with cells expressing Jun-Nt or Fos-Ct. The HTS was performed in solid black 96 well plates. Wells in columns 1-11 were seeded with 50 μL cells from pool A while column 12 received 50 μL of cells from pool B. Next, 50 μL of the small compounds at multiple concentrations were added to the cells (final concentration between 400 μM and 3.2 μM, 1% DMSO). We used a Prestwick Chemical library (Sigma, St. Louis, MO, USA) containing 1120 compounds in 96 well plates (note: currently the commercially available library contains 1280 compounds). In each plate, columns 1 and 12 (positive and negative controls respectively) were treated with DMSO. After 48 h of incubation, the media was substituted by 100 μL of PBS and the fluorescence was measured in a VictorX multiplate reader (Perking Elmer, Waltham, MA, USA). Results obtained from the screen were standardized using the Z-Score, calculated as follows: Z-Score = (x − μ)/δ, where x is the raw signal, μ is the mean signal and δ is the standard deviation of all the compound-containing wells of one plate. The Z-Score indicates how many standard deviations a particular compound is above or below the mean of the plate. Primary hits were identified by calculating a Z-score for each compound and applying hit selection criteria; Z-Score < > 1.5.

Hits were confirmed in secondary screening. This time cells included the pRL-Renilla plasmid to monitor for cell death and protein expression. Additionally, the toxicity of the compounds at the screening concentration was evaluated with Cell-Titer Glo (Promega). In each well, equal amounts of cells transfected with the different plasmids were included.

### Viral infections

VeroE6 cells were infected at an MOI of 0.01 with SARS-CoV-2. Next Ethaverine, Rabeprazole, or Ethynylestradiol were added at the appropriate concentration (see figure 3). Additionally, cells were treated with DMSO. After 48 hrs supernatants were collected and titered by standard plaque assay in VeroE6 cells. Alternatively, MDCKs cells were infected with IAV/WSN/33 at an M.O.I of 0.001. Samples were incubated in the absence or presence of Ethaverine, Rabeprazole, or Ethynylestradiol at the indicated concentrations for 48h. Next, supernatants were collected and titered by the 50 % Tissue Culture Infective Dose (TCID50) method in MDCK cells.

## Acknowledgments

We thank the Generalitat Valenciana (PROMETEO/2019/065) and the Spanish Ministry of Science and Innovation (PID2020-119111GB-I00). We thank P. Selvi for excellent technical assistance. O.Z. is funded by grant PID2020-114546RB-I00 from National Research Agency (Spanish Ministry for Science and Innovation). This publication was also supported by the European Virus Archive GLOBAL (EVA-GLOBAL) project which has received funding from the European Union’s Horizon 2020 research and innovation program under grant agreement 871029.

